# Revisiting Chemoaffinity Theory:Chemotactic Implementation of Topographic Axonal Projection

**DOI:** 10.1101/136291

**Authors:** Honda Naoki

**Author notes:** **Corresponding author:** Honda Naoki Address: Building F, Yoshidakonoe, Sakyo, Kyoto 606-8315, Japan. **Competing Interests:** The author has declared that no competing interests exist.

## Abstract

Neural circuits are wired by chemotactic migration of growth cones guided by extracellular guidance cue gradients. How growth cone chemotaxis builds the macroscopic structure of the neural circuit is a fundamental question in neuroscience. I addressed this issue in the case of the ordered axonal projections called topographic maps in the retinotectal system. In the retina and tectum, the erythropoietin-producing hepatocellular (Eph) receptors and their ligands, the ephrins, are expressed in gradients. According to Sperry’s chemoaffinity theory, gradients in both the source and target areas enable projecting axons to recognize their proper terminals, but how axons chemotactically decode their destinations is largely unknown. To identify the chemotactic mechanism of topographic mapping, I developed a mathematical model of intracellular signaling in the growth cone that focuses on the growth cone’s unique chemotactic property of being attracted or repelled by the same guidance cues in different biological situations. The model presented mechanism by which the retinal growth cone reaches the correct terminal zone in the tectum through alternating chemotactic response between attraction and repulsion around a preferred concentration. The model also provided a unified understanding of the contrasting relationships between receptor expression levels and preferred ligand concentrations in EphA/ephrinA- and EphB/ephrinB-encoded topographic mappings. Thus, this study redefines the chemoaffinity theory in chemotactic terms.

**Author Summary:** This study revisited the chemoaffinity theory for topographic mapping in terms of chemotaxis. According to this theory, the axonal growth cone projects to specific targets based on positional information encoded by chemical gradients in both source and target areas. However, the mechanism by which the chemotactic growth cone recognizes its proper terminal site remains elusive. To unravel this mystery, I mathematically modeled a growth cone exhibiting concentration-dependent attraction and repulsion to chemotactic cues. The model identified a novel growth cone guidance mechanism in topographic mapping, highlighting the importance of the growth cone’s unique ability to alternate between attraction and repulsion. Furthermore, an extension of the model provided possible molecular mechanisms for contrasting two types of topographic mappings observed in the retinotectal system.

## Introduction

During development, neurons extend axon and dendrites [1–3] and axonal growth cones chemotactically migrate in response to extracellular guidance cue gradients and connect to their target sites. Because this axon guidance is a fundamental process in wiring neural circuits, many guidance cues and receptors have been identified and their functional roles (e.g., attraction or repulsion) have been extensively investigated [4–6]. The growth cone’s chemotactic properties are thus being unveiled at the molecular level, but the chemotactic mechanisms of neural circuit construction remain mysterious at the macroscopic level. I addressed this issue by investigating topographic maps, the ordered axonal projections ubiquitous in the sensory nervous system. The best-studied example is in visual system, where retinal ganglion cells (RGCs) project their axons to the optic tectum and/or superior colliculus (SC) while keeping an initial positional relation [7].

The most important concept of topographic map formation is the “chemoaffinity theory” proposed by Roger Sperry in 1940s [8]. Sperry proposed that chemical labels form gradients in source and target areas, allowing a projecting axon to recognize its target site. The theory’s molecular basis was identified with the discovery of gradients of erythropoietin-producing hepatocellular (Eph) receptors and their ligands, ephrins, in the retina (source area) and tectum (target area) [9,10]. Ephs and ephrins are classified into two families, A and B, that encode orthogonal topographic maps in the retina and tectum (**Fig. 1**).

**Figure 1:**
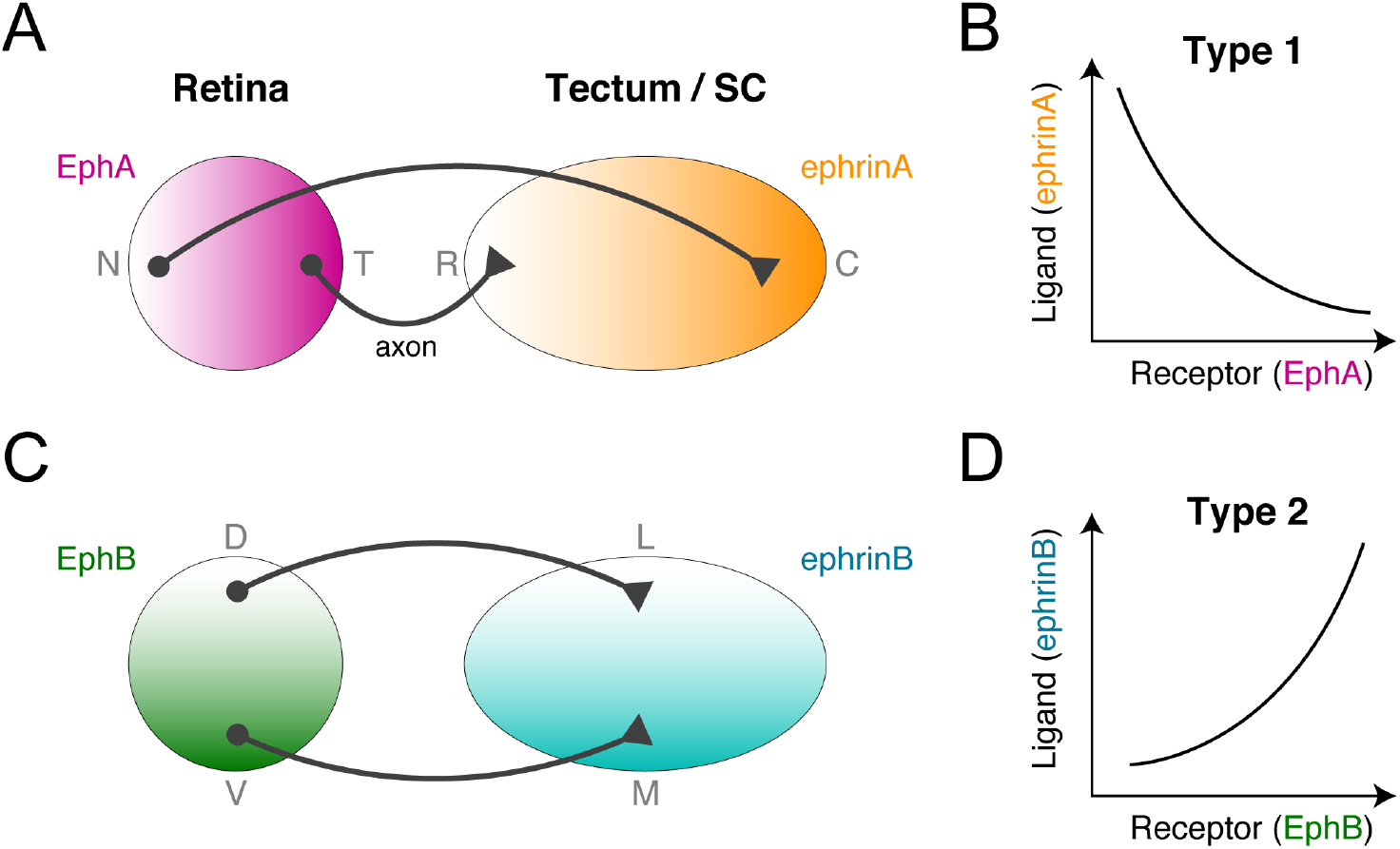
Two types of topographic maps in the retinotectal system. (A, B) Topographic mapping from the retina to the tectum is encoded by orthogonal gradients of EphA and EphB receptors in the retina and of their ligands, ephrinA and ephrinB, in the tectum or SC. (C, D) The EphA/ephrinA- and EphB/ephrinB-encoded topographic mappings exhibit opposite receptor expression level-dependent ligand concentration preferences. These were categorized as types 1 and 2 in this study.

The EphA receptor gradient along the retina’s nasal-temporal axis topographically corresponds to the ephrinA gradient along the tectum’s rostral-caudal axis (**Fig. 1A**). On the orthogonal coordinates, the EphB receptor gradient along the retina’s dorsal-ventral axis corresponds to the ephrinB gradient along the tectum’s medial-lateral axis (**Fig. 1C**). These facts suggest that RGC growth cones chemotactically migrate to their terminal zones guided by ligand concentrations reflective of receptor expression levels. Because ephrinA and ephrinB act as both attractants and repellents in a concentration-dependent manner [11–13], it is possible that growth cones switch between attraction and repulsion around the terminal zone, but the chemotactic mechanism for decoding destination from dual gradients (i.e., receptor and ligand) is unknown.

The EphA/ephrinA- and EphB/ephrinB-encoded topographic maps differ in that the RGCs with higher EphA receptor expression prefer lower tectal ephrinA concentrations (**Fig. 1B**), whereas the RGCs with higher EphB receptor expression prefer higher tectal ephrinB concentrations (**Fig. 1D**). In other words, the retinotectal system’s two kinds of topographic mapping have opposite receptor expression-dependent ligand concentration preferences. How the growth cone’s chemotactic system implements these opposite preferences is also unknown.

Topographic mapping has been extensively investigated with computational models for four decades [14], but all previous models featured growth cones reaching their terminal zones by heuristically-designed chemoaffinity [15–25] While these models provided insights into the outcomes of surgical experiments in the retinotectal system [15–17] and the abnormal maps resulting from misexpression of Ephs or ephrins [15,18–25], none addressed how the intracellular mechanism of growth cone chemotaxis achieves chemoaffinity.

I sought to determine the underlying mechanism of topographic mapping implemented by growth cone chemotaxis. To this end, I focused on the growth cone’s unique chemotactic property of being attracted and repelled by the same guidance cues in different biological environments [26,27]. By mathematically modeling growth cone migration regulated by intracellular signaling, I attempted to demonstrate how the growth cone reaches its terminal zone in the tectum by switching attraction and repulsion around a preferred ligand concentration. Through this model, I redefined Sperry’s chemoaffinity theory in terms of chemotaxis.

## Results

I first studied the projecting growth cone’s preference for a specific ligand concentration associated with the correct terminal zone in the target area. The basic idea is that a growth cone switches between attraction and repulsion around a specific preferred concentration; if the growth cone exhibits attraction and repulsion to lower and higher concentrations, respectively, then it ultimately reaches a location with the preferred concentration. To examine this idea, I mathematically modeled intracellular signaling in chemotactic growth cones.

### Model of bidirectional chemotactic response

The model growth cone was equipped with an intracellular activator (A) and inhibitor (I) of their effector (E), where A and I were upregulated by guidance cues and E regulates the growth cone motility (**Fig. 2A, B**). This activator-inhibitor framework has been commonly observed in both neural and non-neural chemotactic cells [28–31]. For simplicity, a one-dimensional coordinate (*x*) across the growth cone was modeled as {*x*| − *L*/*2* ≤ *x* ≤ *L*/*2*}, where *L* indicates its length. The reaction-diffusion dynamics of A and I were described by

**Figure 2:**
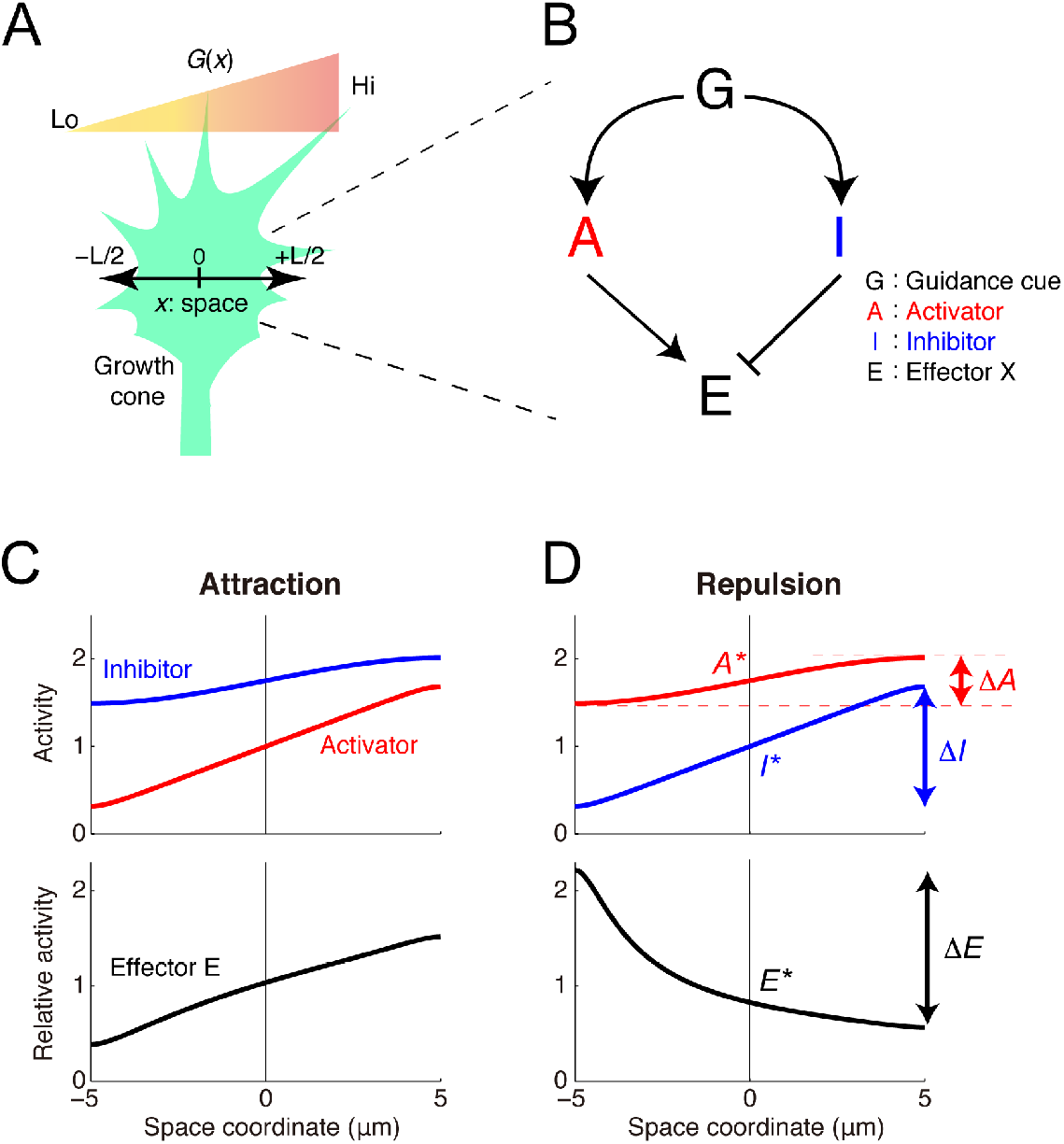
The model of the intracellular growth cone chemotactic process. **(A)** A schematic of the one-dimensional model growth cone encountering an extracellular gradient of guidance cues. **(B)** The model growth cone’s components: a guidance cue (G) regulates an activator (A) and an inhibitor (I) of the effector (E). **(C, D)** Following exposure to a linear extracellular gradient of G (*G*(*x*) = *G*^∗^ + *gx*), gradients of A and I are formed across the growth cone, thereby forming a gradient of E. If the gradient of E orients to the extracellular gradient (Δ*E* > 0), then the growth cone shows attraction (C), but otherwise (Δ*E* < 0), it shows repulsion (D). The model parameters are listed in **Table 1**.

**Table 1:** Parameters.

| Parameter | Unit | Fig. 2C | Fig. 2D | Fig. 3B | Fig. 3C | Fig. 3D | Fig. 3E |
| --- | --- | --- | --- | --- | --- | --- | --- |
| $L$ | $\mu\text{m}$ | 10 | 10 | — | — | — | — |
| $G^*$ | $\mu\text{M}$ | 10 | 10 | — | — | — | — |
| $g$ | $\mu\text{M}/\mu\text{m}$ | 0.75 | 0.75 | — | — | — | — |
| $D_A$ | $\mu\text{m}^2/\text{s}$ | 1 | 100 | 1 | 20 | 1 | 20 |
| $k_A$ | $\text{s}^{-1}$ | 5 | 2 | 20 | 1 | 20 | 1 |
| $c_A$ | $\mu\text{M}/\text{s}$ | 0 | 0 | 150 | 0.05 | 150 | 0.05 |
| $\alpha_A$ | $\text{s}^{-1}$ | 0.5 | 0.35 | 10 | 2.5 | 5 | 5 |
| $D_I$ | $\mu\text{m}^2/\text{s}$ | 100 | 1 | 20 | 1 | 20 | 1 |
| $k_I$ | $\text{s}^{-1}$ | 2 | 5 | 1 | 20 | 1 | 20 |
| $c_I$ | $\mu\text{M}/\text{s}$ | 0 | 0 | 0.05 | 150 | 0.05 | 150 |
| $\alpha_I$ | $\text{s}^{-1}$ | 0.35 | 0.5 | 5 | 10 | 5 | 10 |

Parameter values used in figures 2 and 3 were listed.

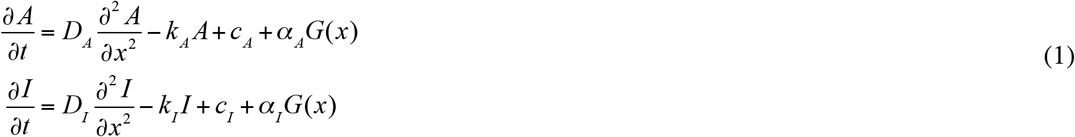

with reflecting boundaries at both ends (*x* = ±*L*/*2*), where *A* and *I* represent the activities of A and I, respectively, *D_Z_*, *k_Z_*, *c_Z_*, and *α_Z_* (*Z* ∈ {*A*, *I*}) denote the diffusion constant, decay rate, constant input, and efficacy, respectively, of the guidance cue’s signal transmission, and *G*(*x*) represents the guidance cue concentration at *x*. The activity of E was determined by the ratio of A’s activity to I’s, i.e., (*x*) = (*x*)/*I* (*x*), which is reasonable if E is regulated by a push-pull enzymatic reaction between A and I [32,33]. The growth cone’s migration was driven by the relative spatial polarity of E as Δ*E*/*E*^∗^, where *ΔE* and *E^*^* indicate the spatial difference of E across the growth cone (i.e., (*L*/*2*) − (−*L*/*2*)) and the baseline activity of E (i.e., (0)), respectively. This property was stated as the Weber-Fechner law, in which the detectable spatial polarity of E varies because of the scale of the concentration of E [34]. Indeed, the Weber-Fechner law has been found in several types of chemotactic cells [35–40]. By analytically solving the model (see Methods), I demonstrated that it produced opposite polarities for *ΔE* depending on the parameters (**Fig. 2C, D**); when Δ*E* > 0, the growth cone was attracted and migrated along the gradient, but when Δ*E* < 0, the growth cone was repelled and turned against the gradient.

### Establishment of preferred concentration by switching attraction and repulsion

I examined how chemotactic responses vary with absolute concentrations in the gradient. My previous study [27] showed that the steady-state response of Δ*E*/*E*^∗^ was presented by

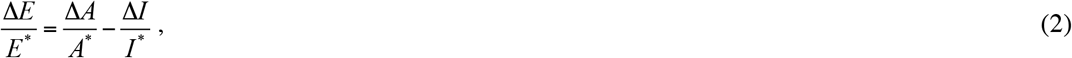

where *A^*^* and *I^*^* denote the baseline activities of A and I, respectively (i.e., *A*^∗^ = *A*(0) and *I*^∗^ = *I*(0)), and Δ*A* and Δ*I* denote the spatial differences of A and I, respectively, across the growth cone (i.e., Δ*A* = *A*(*L*/*2*) − *A*(−*L*/*2*) and Δ*I* = (*L*/*2*) − *I*(−*L*/*2*)) (see **Fig. 2D**). *Z^*^* and *ΔZ* (*Z* ∈ {*A*, *I*}) were analytically derived (see Methods). By substituting these into equation (2), I found that four chemotactic response patterns were generated depending on parameters (**Fig. 3A**): unidirectional repulsion, unidirectional attraction, bidirectional repulsion-to-attraction, and bidirectional attraction-to-repulsion (BAR). In the former two patterns, the growth cone always exhibited attraction or repulsion, meaning that it preferred higher or lower concentrations, respectively (**Fig. 3C, D**). In bidirectional repulsion-to-attraction, the growth cone preferred either higher or lower concentrations depending on the initial concentration (**Fig. 3B**). Finally, in BAR, the growth cone avoided both higher and lower concentrations but preferred a specific concentration by switching attraction and repulsion at that concentration (**Fig. 3E**). I hypothesized that this BAR pattern could play a fundamental role in topographic map formation.

**Figure 3:**
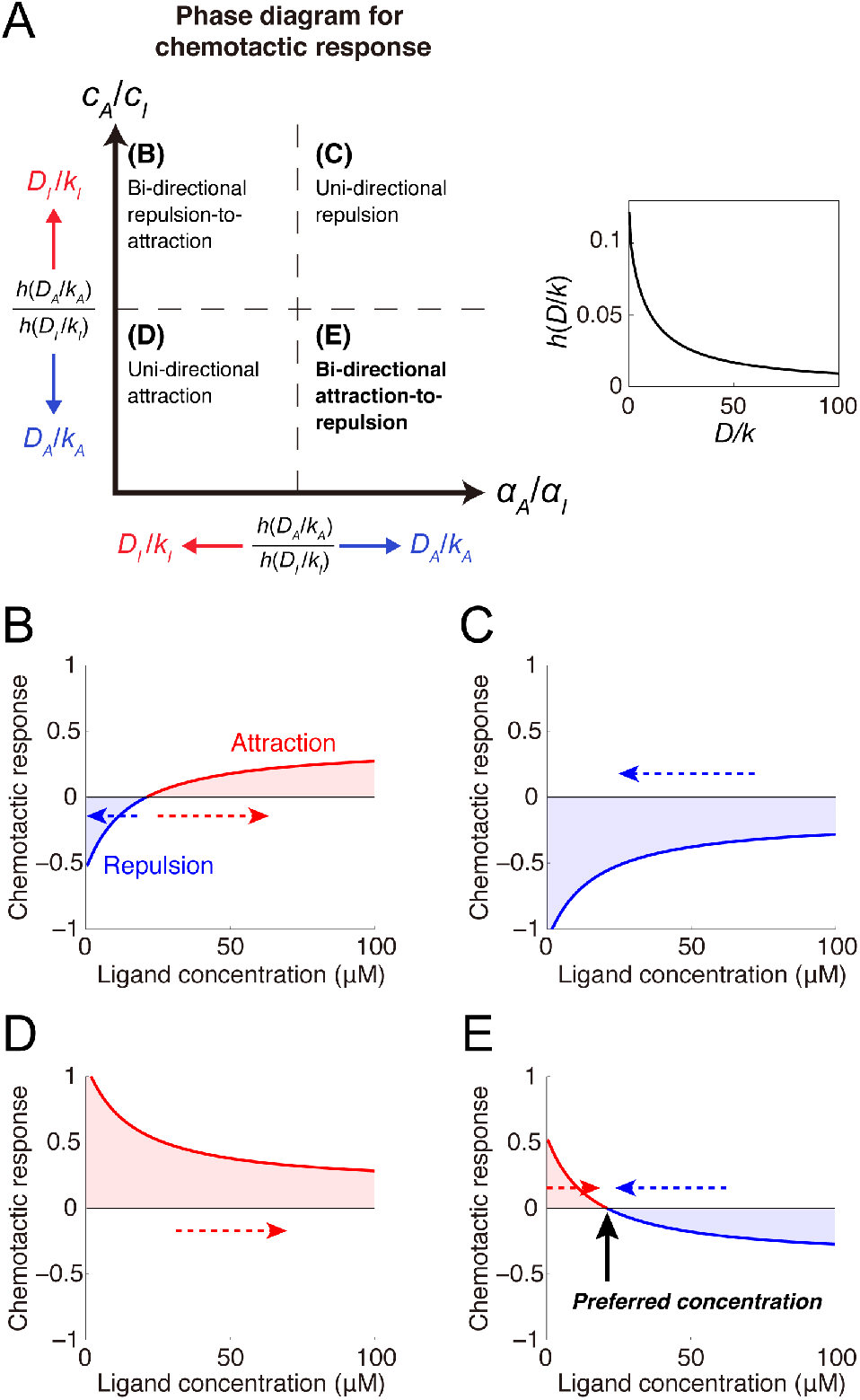
Mechanism of ligand concentration preferences by switching attraction and repulsion. **(A)** Phase diagram depicting parameter regions of the four chemotactic response patterns. The dashed lines indicate critical lines corresponding to a ratio of *h*(*D*_*A*_/*k*_*A*_) to *h*(*D*_*I*_/*k*_*I*_). Because *h*(*D*/*k*) is a monotonically decreasing function of *D*/*k* (inset), the critical lines move with changes in *D*_*A*_/*k*_*A*_ and *D*_*I*_/*k*_*I*_. **(B-E)** Various chemotactic responses (i.e., Δ*E*/*E*^∗^) to guidance cue concentrations were derived: **(B)** bidirectional repulsion-to-attraction, **(C)** unidirectional attraction, **(D)** unidirectional repulsion and **(E)** bidirectional attraction-to-repulsion (BAR). Dashed arrows indicate the direction of concentration changes resulting from attractive or repulsive migration. In the BAR response, the x-intercept indicated by the black arrow corresponds to the preferred guidance cue concentration. The model parameters are listed in **Table 1**.

### Model of topographic mapping

Assuming that the growth cone exhibited the BAR pattern, I studied how receptor expression levels affected the preferred concentration. To this end, the receptor was incorporated into the model as follow:

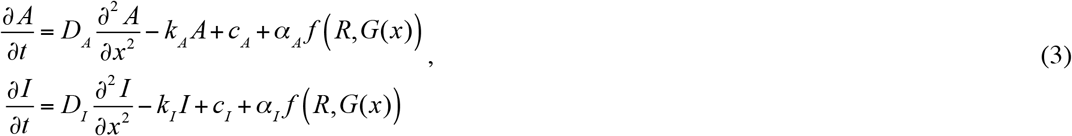

where *R* represents the expressed receptor’s density, and *f* (*R*, *G*) represents the density of the receptor’s active form depending on the guidance cue concentration. By analyzing this model based on equation (2) (see Methods), I found that whether the preferred concentration, *G_pref_*, decreases or increases with *R* was determined by the sign of derivatives of *f* (*R*, *G*) with respect to *R* and *G*:

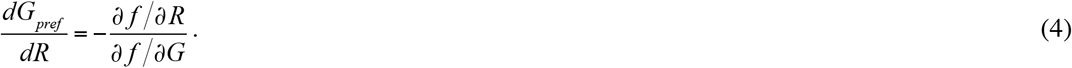

Therefore, *f* (*R*, *G*), which represents how the guidance cue signal is transmitted to A and I through the receptor, is a crucial factor in the receptor expression level-dependent preferred ligand concentration. I next studied specific examples of (*R*, *G*).

### Type 1 mapping encoded by EphA/ephrinA

I considered a scenario in which the receptors were activated by guidance cue binding (**Fig. 4A**), which is described by *f* (*R*, *G*) = *RG*/(*K* + *G*), where *K* is the dissociation constant of binding reaction between the receptor and guidance cue (i.e., a ratio of unbinding rate to binding rate). I then calculated the preferred concentration based on equation (2) and found that it decreased with the receptor expression level (**Fig. 4B**) as

**Figure 4:**
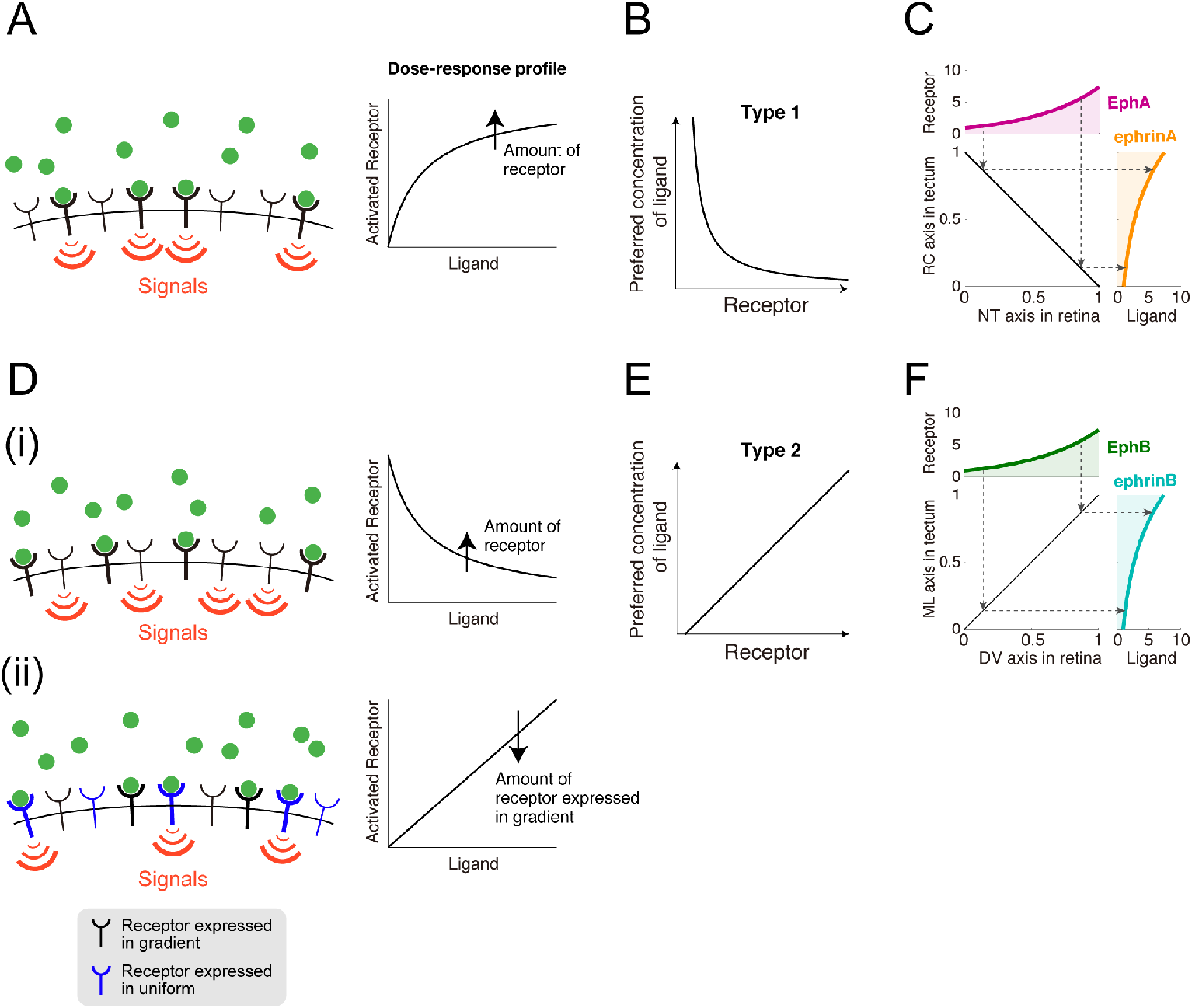
Topographic mapping implemented by growth cone chemotaxis. **(A)** The receptors were activated by guidance cue binding. Dose-response of receptor activation was plotted in right panel. **(B)** The chemotactic growth cone in (A) prefers a specific ligand concentration that is inversely proportional to the receptor expression level. **(C)** Linear topographic mapping was produced by an EphA gradient along the retinal nasal-temporal (NT) axis and an ephrinA gradient along the tectal rostral-caudal (RC) axis. Dashed arrows indicate corresponding receptor expression levels and preferred ligand concentrations. The applied gradients were *R_NT_*(*x_NT_*) = *R_NT_o__exp*(*q_A_*(*x_NT_*/*σ_NT_*)) and 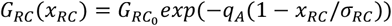. **(D)** Two possible molecular mechanisms by which the guidance cue is transduced to an intracellular signal through the receptor. **(D(i))** Guidance cue-unbound receptors were active. **(D(ii))** Two kinds of receptors competitively bind the guidance cue so that these receptors effectively suppress each other. **(E)** The chemotactic growth cone in (D) prefers a specific ligand concentration that linearly increases with the receptor expression level. **(F)** Linear topographic mapping was produced by an EphB gradient along the retinal dorsal-ventral (DV) axis and an ephrinB gradient along the tectal medial-lateral (ML) axis. The applied gradients were 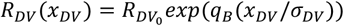and 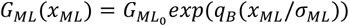.

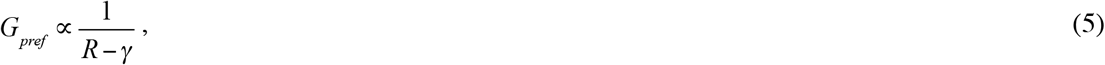

where *γ* is a positive constant determined by the model parameters. This is consistent with type 1 topographic mapping in which higher EphA levels result in the growth cone preferring smaller ephrinA concentrations (**Fig. 1a, b**). If the receptor expression level is greater than *γ*, this relationship produces a linearly ordered topographic map with exponential distributions of retinal EphA and tectal ephrinA (**Fig. 4C**).

### Type 2 mapping encoded by EphB/ephrinB

For the mechanism of type 2 EphB/ephrinB-encoded topographic mapping, I tested two biologically plausible hypothetical *f* (*R*, *G*) expressions. First, guidance cue-unbound receptors might trigger intracellular signaling, which can be expressed by *f* (*G*) = *RK*/(*K* + *G*) (**Fig. 4D(i)**) (see Discussion for its biological relevance). For this hypothesis, I found that the preferred concentration increases with the receptor expression level (**Fig. 4E**) as

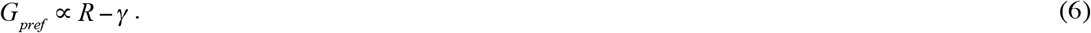

This is consistent with the fact that higher EphB levels result in the growth cone preferring higher ephrinB concentrations (**Fig. 1C, D**) (see Methods). This linear relationship (equation (6)) produced a linearly ordered topographic map with exponential distributions of retinal EphB and tectal ephrinB (**Fig. 4F**).

In the second hypothesis, I assumed that two kinds of receptor competitively bind the limited ligands (**Fig. 4D(ii)**) (see Discussion for its biological relevance). One kind is uniformly expressed across the retina and the guidance cue-bound form triggers intracellular signaling. The other is expressed in gradients across the retina and indirectly inhibits the uniformly expressed receptor by competitively binding the ligand. This case is described by *f* (*R*, *G*) = *R*_c_/(*K* + *R*_c_ + *R*) (Methods), where *R* and *R_c_* indicate densities of the receptors expressed in gradients and uniformly, respectively, and *K* indicates the dissociation constant of the receptor and ligand. For this hypothesis, I also found that the preferred concentration increases with the receptor expression level as

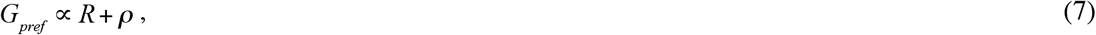

where *ρ* is a positive constant determined by the model parameters. Thus, this hypothesis also explained the type 2 EphB level-dependent preferred ephrinB concentration.

## Discussion

I presented a mathematical model of chemotactic response of the growth cone to reveal how topographic map is formed by the growth cone chemotaxis. In my model, for the sake of simplicity, I assumed that the migration direction of the growing axon was determined by polarity of the growth cone signaling. The real mechanism must be more complicated than what assumed in my model. However, the minimalist model I developed was very informative and provided a novel chemotaxis-based logic of chemoaffinity theory for topographic mapping. I demonstrated that the model could generate both attractive and repulsive responses depending on absolute concentrations along the gradient. Such bidirectionality endows the growth cone with the preference for a specific guidance cue concentration by switching between attraction and repulsion around that concentration. I also determined the conditions of EphA/ephrinA- and EphB/ephrinB-encoded topographic mapping, in which the preferred concentration decreases and increases, respectively, with the receptor expression level. This study therefore redefined Sperry’s chemoaffinity theory in terms of chemotaxis.

### Ephrins as attractants and repellents

If ephrinA is a repellent, as classically thought [41], then all RGC growth cones must project to the tectum’s rostral end, which has the lowest ephrinA concentration. However, this is not the case; even without tectal space competition between projecting axons, the RGC axons project to the correct terminal zone in the tectum [42]. This contradiction can be resolved simply by regarding ephrinA as both an attractant and a repellent. In fact, ephrinA has been reported to be an attractant or a repellent in a concentration-dependent manner [11]. EphrinB has been regarded as both an attractant and a repellent [12,13]. However, their underlying mechanism was largely unknown. In this study, I demonstrated how ephrinA and ephrinB could indeed work as both attractants and repellents for the chemotactic growth cone.

### Signal transmission through EphA and EphB

I demonstrated that whether the preferred ligand concentration decreases or increases with the receptor expression level is determined by whether the guidance cue and the receptor positively or negatively affect intracellular signaling (equation (4)). As the mechanism of EphA/ephrinA-encoded type 1 topographic mapping, I reasonably assumed that ephrinA-bound EphAs trigger intracellular signaling (**Fig. 4A**), but for type 2 topographic mapping, I tested two hypothetical EphB/ephrinB regulation schemes. The first hypothesis was that ephrinB-unbound EphBs, rather than bound ones, trigger intracellular signaling (**Fig. 4D(i)**). This seems inconsistent with a property of tyrosine kinase-type receptors, which are activated by ligand binding through phosphorylation [43,44], but it has recently been reported that Ephs can be ligand-independently activated by hemophilic Eph-Eph interactions [45], suggesting that ephrinB-bound and - unbound EphBs could generate different signals. The first hypothesis was thus biologically feasible, but further experimental investigation is needed. The second hypothesis was that two kinds of receptor, which are expressed uniformly or in gradients across the retina, competitively bind the ligand (**Fig. 4D(ii)**). This fits the expression profiles of EphB subtypes in the chicken retina well; EphB2 and EphB3 are expressed in gradients across the retina, whereas EphB1 is uniformed expressed [7]. My hypothesis thus offers experimentally testable predictions concerning EphB/ephrinB regulation.

### Functional difference between two types of topographic mappings

It is worth mentioning functional difference between type 1 and type 2 of topographic mappings. I deduced that local accuracy of axonal projection is determined by multiplication of three factors: 1. spatial derivatives of receptor expression profile in retina (upper panels in **Fig. 4C and F**), 2. steepness of mapping function from receptor expression to preferred ligand concentration (**Fig. 4B and E**) and 3. spatial derivatives of ligand in tectum (right panels in **Fig. 4C and F**). In type 1 topographic mapping, while multiplication of the first and third factors, i.e., (*dR_NT_*/d*x_NT_*)(*dG_RC_*/d*x_RC_*), is constant (**Fig. 4C**), the second factor, i.e., the steepness of mapping function, increases as EphA expression decreases (**Fig. 4B**). On the other hand, in type 2 topographic mapping, while the second factor, i.e., the steepness of mapping function, is constant (**Fig. 4E**), multiplication of the first and third factors, i.e., (*dR_DV_*/d*x_DV_*)(*dG_ML_*/d*x_ML_*), increases with EphB expression. Thus, it can be predicted that axonal projection from nasal ventral retinal region associated with lower EphA and higher EphB expression could be more precise than that other retinal region.

### Species-dependent pattern of axonal projections

RGCs’ axonal projection patterns in the optic tectum or SC are species-dependent. In higher vertebrates (i.e., mammals and birds), the axons overshoot their terminal zones and subsequently form branches [7], while in lower vertebrates (i.e., fish and amphibians), the growth cones directly reach and stop in their terminal zones [7] despite being initially misrouted [46]. The latter case suggests that the chemotactic system implements chemoaffinity, which I investigated as the mechanism of topographic mapping. The growth cone’s chemotaxis might therefore play a fundamental role in topographic mapping, while axonal overshoot and branching might facilitate exploration of the terminal zone. My model could understand the axonal overshoot by incorporating transient dynamics of activator and inhibitor, instead of steady state assumption. On the other hand, how the axon generates branches is out of scope of my model.

### Comparison with previous chemotaxis models

Chemotactic gradient sensing has been computationally studied mainly for non-neural chemotactic cells [40,47–51] like *Dictyostelium discoideum* and immune cells, though attraction to guidance cues has been only paid attention. On the other hand, there are a couple of computational models for the growth cone chemotaxis alternating attraction and repulsion [27,52]. These models, whether applied to neural or non-neural cells, primarily addressed intracellular signaling consisting of activators and inhibitors. In the non-neural cells, the activator and inhibitor were thought to be PI3K and PTEN [28], respectively, or RasGEF and RasGAP [29], respectively. In the growth cone, CaMKII and PP1 were thought to work as the activator and inhibitor, respectively [27,30,31,52], which regulate cellular motility via Rho GTPases [53]. In short, chemotactic responses could be understood from the activator-inhibitor framework [54], so I hypothesized that RGC chemotaxis is also regulated by an activator-inhibitor system, although the intracellular signaling pathway of Eph/ephrin has not been fully identified.

### Comparison with previous models of topographic mapping

There have been many computational studies on topographic mapping [14]. These studies did not focus on the intracellular mechanism of growth cone chemotaxis, but instead developed models with heuristically designed chemoaffinity (e.g., optimization of energy function) by which the growth cone reaches its correct terminal zone. Given such chemoaffinity, these models potentially gave insights into more system-level phenomena, such as abnormal maps resulting from surgical experiments in the retinotectal system [15–17] and from the misexpression of Eph or ephrin [15,18–25]. These models included several factors not included in my model, such as axon competition for tectal space [55] and counter-gradients of Ephs and ephrins in the retina and tectum [56]. Several models have also addressed a question of how synaptic connection is refined by activity-dependent synaptic plasticity mechanism after activity-independent axon guidance [20,57–59]. Therefore, I must stress that my model does not compete with previous models, but rather can explain the underlying mechanism by which growth cones can chemotactically implement the previous models’ heuristically designed chemoaffinity.

## Methods

### Theory for chemotactic response

Suppose a shallow extracellular gradient because growth cones are known to detect few percent difference of concentrations across the growth cone [60–64]. I then assumed that the intracellular gradients of A and I, *A*(*x*) and *I*(*x*), were shallow and slightly perturbed from their activities at *x* = 0. The activity of E at *x* could be linearized as

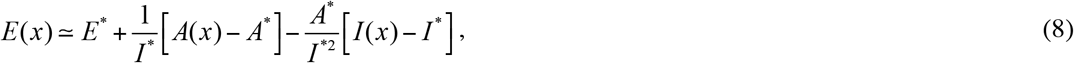

where *A*^∗^ = *A*(0), *I*^∗^ = *I*(0), and *E*^∗^ = *E*(0) = *A*^∗^/*I*^∗^. The relative spatial difference of E across the growth cone was calculated by

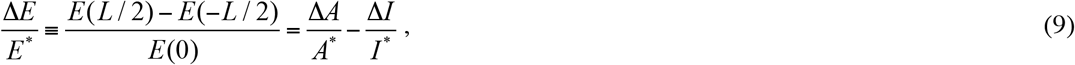

where Δ*A* and Δ*I* indicate the spatial differences of A and I, respectively, across the growth cone.

### Distribution of A and I

For both A and I, I calculated the intracellular distribution exposure to an extracellular gradient, *G*(*x*). Green’s function of ∂*Z*/∂𝑡 = *D*_*z*_ (∂^2^*Z*/∂*x*^2^) – *k*_Z_ Z was analytically derived using the method of separation of variables:

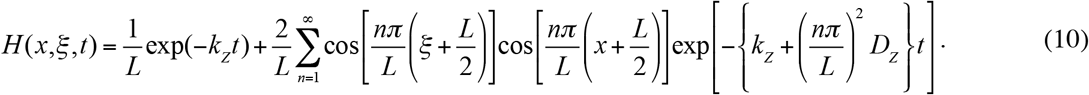

A steady-state solution of equation (3) was thus obtained by

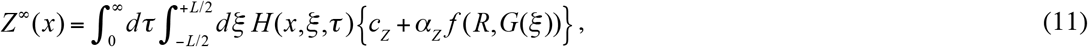

where *Z* represents either *A* or *I*. Note that *f* (*R*, *G*(*x*)) = *G*(*x*) in equation (1). Because the growth cone is so small that *G*(*x*) could be modelled as a shallow linear gradient, (*R*, (*x*)) can be linearized by *f* (*R*, *G*^∗^) +*gx*, where *G*^∗^ = (0) and *g* = (∂*f*/∂*G*|_G=G_∗)(𝑑*G*/𝑑*x*|_x=0_). This led to

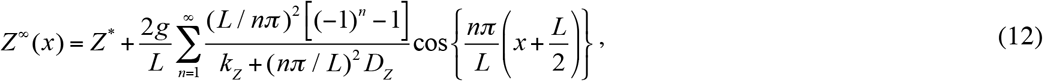

where *Z^*^* indicates baseline activity, i.e., *Z*^∗^ = *Z*^*∞*^(0):

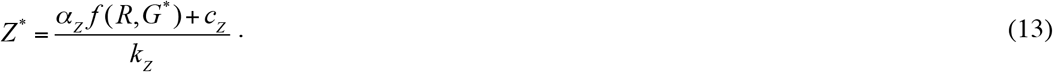

By numerical simulation of the reaction-diffusion dynamics, I confirmed that equation (12) was exact. The spatial difference of *Z* then becomes

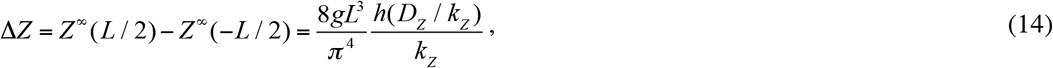

where

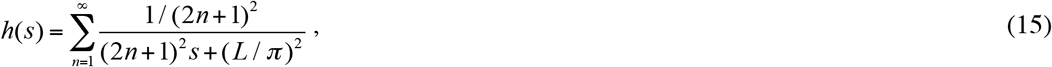

which is a monotonically decreasing function converging to 0 (inset of **Fig. 3A**).

### Conditions for four chemotactic response patterns

I calculated the growth cone’s concentration-dependent chemotactic responses. By substituting *Z*^*^ as described by equation (13) for *A*^*^ and *I*^*^ in equation (2) and substituting *ΔZ* as described by equation (14) for *ΔA* and *ΔI* in equation (2), I obtained

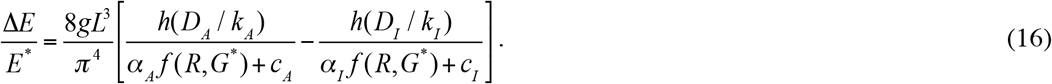

Equation (16) exhibits four response patterns to *G^*^*: all positive, all negative, negative-to-positive, and positive-to-negative, which correspond to unidirectional attraction, unidirectional repulsion, bidirectional repulsion-to-attraction, and BAR, respectively (**Fig. 3B-E**). The response patterns’ parameter regions were derived under the condition of ∂*f* /∂*G* > 0 (**Fig. 3A**). For example, the BAR response pattern is characterized by attraction at lower concentrations (i.e., *E*/*E*^∗^|_G_∗_=0!_ > 0) and repulsion at *G*^∗^ = *∞* (i.e., which leads to

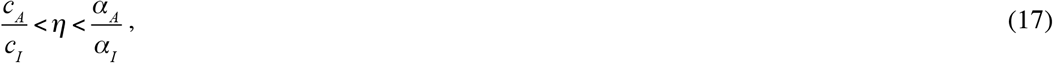

where 𝜂 = *h*(*D*_A_/*k*_A_)/*h*(*D*_I_/*k*_I_).

### Preferred concentration in the BAR response pattern

Growth cones with the BAR response pattern prefer a specific concentration of *G^*^* at which Δ*E*/*E*^∗^ = 0. In the equation (3) model, setting Δ*E*/*E*^∗^ = 0 in equation (16) leads to

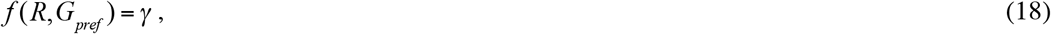

where 𝛾 = (𝜂𝑐_*I*_ – 𝑐_*A*_)/(𝛼_*A*_ – 𝜂𝛼_I_). The preferred concentration with a specific *f* (*R*, *G*) can be calculated with equation (18). In the equation (1) model, *f* (*R*, *G*) = *G*, thus *G*_*pref*_= 𝛾. If *f* (*R*, *G*) = *RG*/(*K* + *G*), *G*_*pref*_ = 𝛾*K*/(*R* − 𝛾) (equation (5); **Fig. 4A**). If *f* (*R*, *G*) = *RK*/(*K* + *G*), *G* _*pref*_= (*K*/𝛾)(*R* − 𝛾) (equation (6); **Fig. 4D(i)**). If *f* (*R*, *G*) = *R*_c_ /(*K* + *R*_c_ + *R*), *G*_*pref*_ = (𝛾/*R*_c_)/(*R* + *K* + *R*_c_) (equation (7); **Fig. 4D(ii)**). Total differentiation of equation (18) leads to (∂*f*/∂*R*) + (∂*f*/∂*G*_*pref*_) _*pref*_ = 0, which in turn leads to

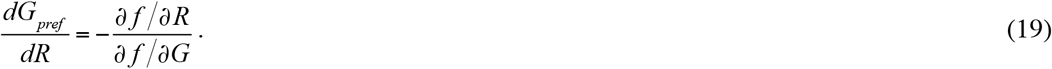

### Competitive binding of limited ligands by two receptors

I assumed a scenario in which two kinds of RGC-expressed receptors competitively bind limited ligands with identical kinetics. Note that the two assumed kinds are expressed either uniformly or in gradients across the retina. Such dynamics are described by

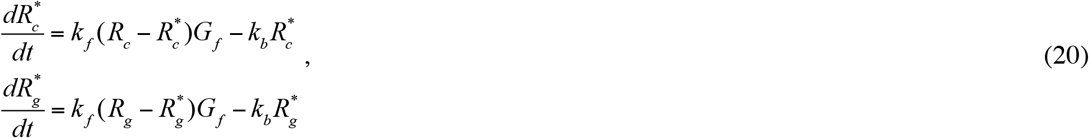

where R_*j*_, 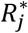, and *G_f_* (𝑗 ∈ {𝑐, *g*}) indicate densities of the total receptors, guidance cue-bound receptors, and free guidance cues, respectively, and *k_f_* and *k_b_* indicate forward and backward reaction rates, respectively. The total guidance cue concentration is conserved as 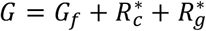 At steady state, 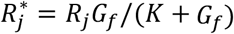 where 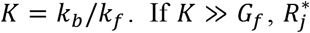 can be approximated as (*R*_j_/*K*)*G*_*f*_, and the steady state of 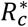 depending on *G* is then described by

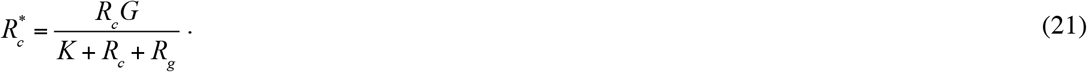

## Funding

This study was partially supported by the Platform Project for Supporting in Drug Discovery and Life Science Research (Platform for Dynamic Approaches to Living System) from Japan Agency for Medical Research and Development (AMED) and Grant-in-Aids for Young Scientists (B) (No. 25730177 and No. 16K16147) from the Ministry of Education, Culture, Sports, Science and Technology (MEXT), Japan. The funders had no role in study design, data collection and analysis, decision to publish, or preparation of the manuscript.

## Acknowledgements

I am grateful to Dr. Michiyuki Matsuda for his valuable comment and financial assistance. I also thank Drs. Makoto Nishiyama, Kyonsoo Hong and Shin Ishii for their stimulating discussions, Drs. Yohei Kondo and Masataka Yamao for critically reviewing the manuscript and Akiko Kawagishi for her technical assistance.

